# Gut mélange à trois: fluctuating selection modulated by microbiota, host immune system, and antibiotics

**DOI:** 10.1101/2021.12.30.474527

**Authors:** Hugo C. Barreto, Beatriz Abreu, Isabel Gordo

## Abstract

Iron is critical in host-microbe interactions, and its availability is tightly regulated in the mammalian gut. Antibiotics and inflammation can perturb iron availability in the gut, which could alter host-microbe interactions. Here, we show that an adaptive allele of *iscR*, a major regulator of iron homeostasis of *Escherichia coli*, is under fluctuating selection in the mouse gut. *In vivo* competitions in immune-competent, immune-compromised, and germ-free mice reveal that the selective pressure on an *iscR* mutant *E. coli* is modulated by the presence of antibiotics, the microbiota, and the immune system. *In vitro* assays show that iron availability is an important mediator of the *iscR* allele fitness benefits or costs. We identify Lipocalin-2, a host’s immune protein that prevents bacterial iron acquisition, as a major host mechanism underlying fluctuating selection of *iscR*. Our results provide a remarkable example of strong fluctuating selection acting on bacterial iron regulation in the mammalian gut.

## INTRODUCTION

Iron is one of the most abundant metals on Earth and essential to the majority of life forms, including microorganisms (Schaible and Kaufmann, 2004). The mammalian gut is a melting pot of interactions required for the proper functioning of the host-microbe duet (Nairz et al., 2010; Yilmaz and Li, 2018; Seyoum et al., 2021). In the dynamic environment of the gut, which is the organ that comprises the largest microbial populations in mammals (Whitman et al., 1998), iron availability is tightly regulated and critical for maintaining a healthy status (Zimmermann and Hurrell, 2007). Some studies have shown that the gut microbiota can affect host iron regulation, and vice-versa. For example, rats devoid of microbiota (germ-free) are deficient at retaining and absorbing iron (Reddy et al., 1972), and, consistently, intestinal cells of germ-free mice were also shown to be depleted of iron (Deschemin et al., 2016). As well, *E. coli*, a member of the gut microbiota, increases the expression of ferric iron uptake pathways in the mucus of germ-free mice (Li et al., 2015). Furthermore, gut microbial metabolites can regulate the host’s iron homeostasis (Das et al., 2020) and the host is able to modify bacterial iron uptake through the secretion of Lipocalin-2 (Lu et al., 2019). Lipocalin-2, also known as neutrophil gelatinase-associated lipocalin (NGAL), is an innate immune protein usually found at low levels in healthy hosts (Nielsen et al., 1996; Chassaing et al., 2012) and critical for the maintenance of iron and gut homeostasis (Srinivasan et al., 2012; Singh et al., 2016). Lipocalin-2 is able to limit bacterial iron uptake by binding to siderophores (Goetz et al., 2002), which are iron-scavenging molecules produced by bacteria (Guerinot, 1994). This is particularly important for the host’s ability to control the emergence of pathogens, as mice lacking Lipocalin-2 are more prone to infections (Flo et al., 2004).

Iron intake is also known to affect the gut microbiota. For instance, an association between iron deficiency and low abundance of *Lactobacillus acidophilus* has been described in humans (Balamurugan et al., 2010), and iron-deprived rodents show an increased abundance of *Lactobacillus, Enterobacteriaceae*, and *Enterococcus*, as well as a decreased abundance of *Bacteroides* and *Roseburia*. (Tompkins et al., 2001; Dostal et al., 2012). Furthermore, supplementation with iron in humans and mice is associated with a decrease in abundance of beneficial microbes, such as *Bifidobacterium* and *Firmicutes*, and an increase in *Bacteroides* and *Enterobacteriaceae*, such as *E. coli* (Mevissen-Verhage et al., 1985; Zimmermann et al., 2010; Jaeggi et al., 2015; Constante et al., 2017). Finally, genetic modification of the host’s iron metabolism was shown to change gut microbiota composition and to decrease the abundance of lactic acid bacteria in mice (Buhnik-Rosenblau et al., 2012).

Antibiotic treatment and inflammation can also modulate host iron homeostasis and alter the gut microbiota composition. Following antibiotic treatment, the absorption of iron was shown to be depleted in rabbits (Stern et al., 1954) and rats (Forrester et al., 1962). Antibiotic treatment is well known to cause gut microbiota dysbiosis, and the recovery to the original state, when observed, requires large time periods and depends on several factors, such as the host diet, community context, and environmental reservoirs (Ng et al., 2019). Inflammation can also cause dysbiosis of the gut microbiota (Lupp et al., 2007) and is accompanied by an increase in the levels of fecal Lipocalin-2 (Chassaing et al., 2012; Lu et al., 2019). Accordingly, in inflammatory conditions the host reduces the availability of iron in the gut (Kortman et al., 2014), which ultimately can lead to anemia of inflammation (Wessling-Resnick, 2010; Ganz and Nemeth, 2015). Perturbations caused by antibiotic treatment and inflammatory factors, which change the interplay between host and bacterial iron homeostasis, can thus potentially lead to fluctuating selective pressures that gut bacteria must cope with.

Most studies on how bacterial populations respond to fluctuating environments have been performed under laboratory conditions (Tagkopoulos et al., 2008; Beaumont et al., 2009; Sandberg et al., 2017; Turner et al., 2020; Nguyen et al., 2021). However, naturally fluctuating environments such as those in the mammalian gut are still poorly studied (Nguyen et al., 2020). Theory predicts that when an organism experiences multiple generations in a seasonal environment, fluctuating selection can lead to maintaining polymorphism (Melbinger and Vergassola, 2015; Sæther and Engen, 2015; Novak and Barton, 2017; Erickson et al., 2017). Since several gut commensals have generation times that can be shorter than fluctuations inside their hosts, *e.g*. due to the circadian rhythm, fluctuating selection could contribute to shaping intra-species genetic diversity within hosts.

In *E. coli*, a common member of the mammalian gut microbiota, one of the major regulators of iron homeostasis is the iron-sulfur cluster regulator IscR. IscR is a transcriptional regulator that harbors its own iron-sulfur cluster (Schwartz et al., 2001), which is essential for its function as a sensor of the cellular iron-sulfur cluster pool (Giel et al., 2013; Santos et al., 2015; Esquilin-Lebron et al., 2021). Given the widespread distribution of IscR homologs in bacteria, its importance in host-microbe interactions has been intensively studied (Kim et al., 2009; Lim and Choi, 2014; Miller et al., 2014; Fuangthong et al., 2015). IscR contains three cysteine residues that are required for the coordination of the IscR iron-sulfur cluster (Yeo et al., 2006). In a recent study, mutations in these cysteine residues of IscR emerged when *E. coli* evolved in the inflamed gut of aging mice under antibiotic treatment (Barreto et al., 2020a). The temporal pattern of the *iscR* mutant frequencies showed a strong fluctuation, very distinct from other metabolic mutations which spread to fixation. Here, we test the hypothesis that the iron-sulfur cluster regulator IscR is under fluctuating selection in the mammalian gut, and perform *in vivo* and *in vitro* experiments to dissect the underlying mechanisms contributing to that form of natural selection. Using an *iscR* mutant that is unable to assemble its own iron-sulfur cluster, we demonstrate that fluctuating selection on *iscR* depends on the presence of microbiota and antibiotic treatment and its period on the host adaptive immune system. Through *in vitro* manipulation of iron availability, through either Lipocalin-2 or the iron chelator Dipyridyl, we show that drastic changes in the competitive fitness of the *iscR* mutant should occur in an environment where iron fluctuates, providing a mechanistic explanation for the patterns observed *in vivo*. Consistent with this prediction, a correlation between the abundance of the *iscR* mutant and fecal Lipocalin-2 levels is observed in the gut. In summary, our work provides evidence for biotic and abiotic mechanisms underlying fluctuating selection on a key bacterial process, iron homeostasis in the mammalian gut.

## RESULTS

### *iscR* mutation C92S is under fluctuating selection in the aging gut

IscR senses the cellular iron-sulfur cluster pool via its iron-sulfur cluster, and the occupancy of this cluster is critical to IscR regulation (Santos et al., 2015). The coordination of the IscR iron-sulfur cluster requires the cysteine residues at positions 92, 98, and 104, as well as the histidine residue at position 107 (Yeo et al., 2006). To determine the level of conservation of these residues we compared IscR homologs, obtained from the NCBI non-redundant protein database, using Consensus Finder (Jones et al., 2020). We found these residues to be highly conserved, with a conservation level of 0.97 for cysteine 92, 0.96 for cysteine 98, 0.97 for cysteine 104, and 0.95 for histidine 107 (Figure 1A and Table S1). Consistent with their high conservation, these residues are important for IscR function, as the replacement of these residues with either alanine or serine leads to IscR loss of coordination of its iron-sulfur cluster and locks IscR in a clusterless form (Nesbit et al., 2009; Santos et al., 2014). Amino acid substitutions in one of these residues, specifically C92S, have however been detected during evolutionary adaptation of *E. coli* colonizing the gut of aging mice (Barreto et al., 2020a). Since this mutation should lead IscR to be locked in its clusterless form, it likely changed the regulation of *E. coli* iron homeostasis (Santos et al., 2014). We, therefore, hypothesized that the *iscR* C92S mutant is under fluctuating selection in the gut where iron concentrations can rapidly change (Yilmaz and Li, 2018).

**Figure 1.**
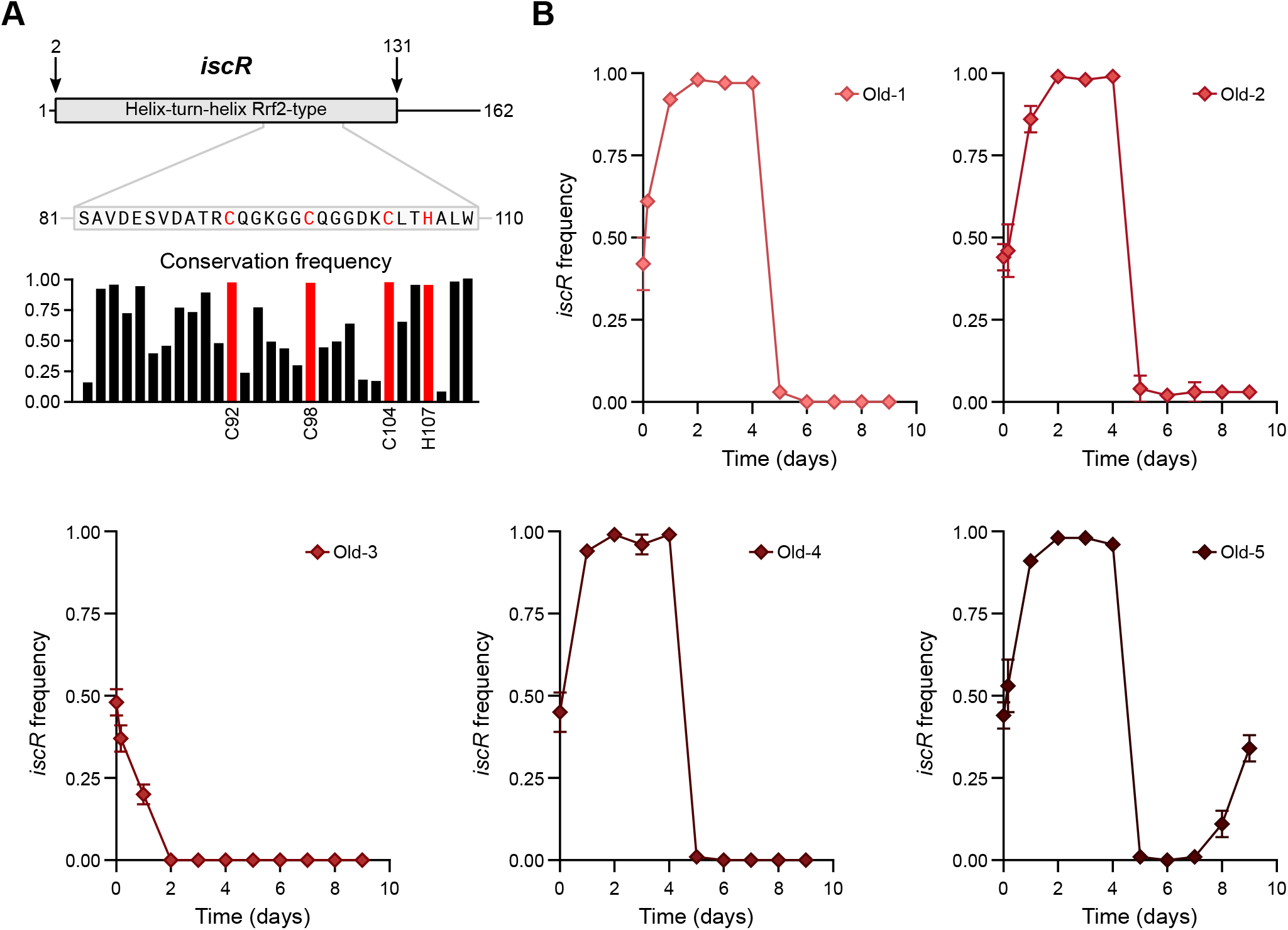
The *iscR* mutation C92S is under fluctuating selection in the aging gut. **(A)** Graphical representation of IscR domains (top), amino acid sequence (middle), and amino acid conservation frequency according to Consensus Finder (bottom). In red are the amino acids that are required for the coordination of the IscR iron-sulfur cluster. **(B)** Temporal dynamics of the YFP-labelled streptomycin-resistant *iscR* mutant when competing against the CFP-labelled streptomycin-resistant wild-type *E. coli* in Old (n = 5) mice that undergo a continuous streptomycin treatment in their drinking water. In panel B the error bars represent the ± 2*SEM. See also Table S1 and S2.

We first tested our hypothesis in old mice treated with the antibiotic streptomycin, the same environment where the *iscR* C92S mutation originally emerged (Barreto et al., 2020a). We colonized 19-month old mice with a 1:1 mixture of the *iscR* C92S mutant, thereafter named *iscR*, (labeled with the yellow fluorescent protein YFP and carrying a streptomycin resistance mutation K43R) and the wild-type *E. coli* (labeled with the cyan fluorescent protein CFP and carrying a streptomycin resistance mutation K43R). We then followed the frequency of the mutant clones throughout time to search for a signal of fluctuating selection. Consistent with our hypothesis, in 4 out of 5 mice, *iscR* is strongly selected for in the first two days (0.13 ± 0.01 SEM per generation), reaching very high frequencies by day four (Figure 1B), after which it is selected against, as its frequency sharply decreases (Figure 1B). The strong change in frequency in the majority of the mice is indicative of a change in the selective pressure on the *iscR* mutant, and with the occurrence of fluctuating selection in the aging gut.

### Fluctuating selection is modulated by the adaptive immune system

Aging is a multifactorial process that affects many physiological traits of the host, such as its microbiota composition, the evolution of its gut commensals, and the functioning of its immune system (Smith et al., 2017; Thevaranjan et al., 2017; Barreto et al., 2020a). Thus, the temporal dynamics observed in the *iscR* frequency could be driven by several factors in the environment of the aging host. We, therefore, asked if the dynamics of *iscR* competing against the wild-type *E. coli* would be different in young adult mice (WT). We observed that, in most young adult mice, *iscR* is highly selected in the first days of the competition reaching very high frequencies after two days (Figure 2A). Similar to the observations in Old mice, an inversion of selection for *iscR* could be detected in all the mice after eight days of competition (Figure 2A). Surprisingly, the timing at which we observed the sharp decrease in *iscR* frequency occurred was shorter in Old than in WT mice (p = 0.003, Unpaired t-test with Welch’s correction), with a mean of 5 days in Old and 10 days in WT mice (Figures 1B and 2A). Remarkably, in 3 out of 6 mice a change in the direction of the selection pressure on *iscR* was observed twice, and in one mouse (WT-2) the selection changed at three instances (Figure 2A). These results demonstrate that *iscR* can also undergo fluctuating selection in young adult mice.

**Figure 2.**
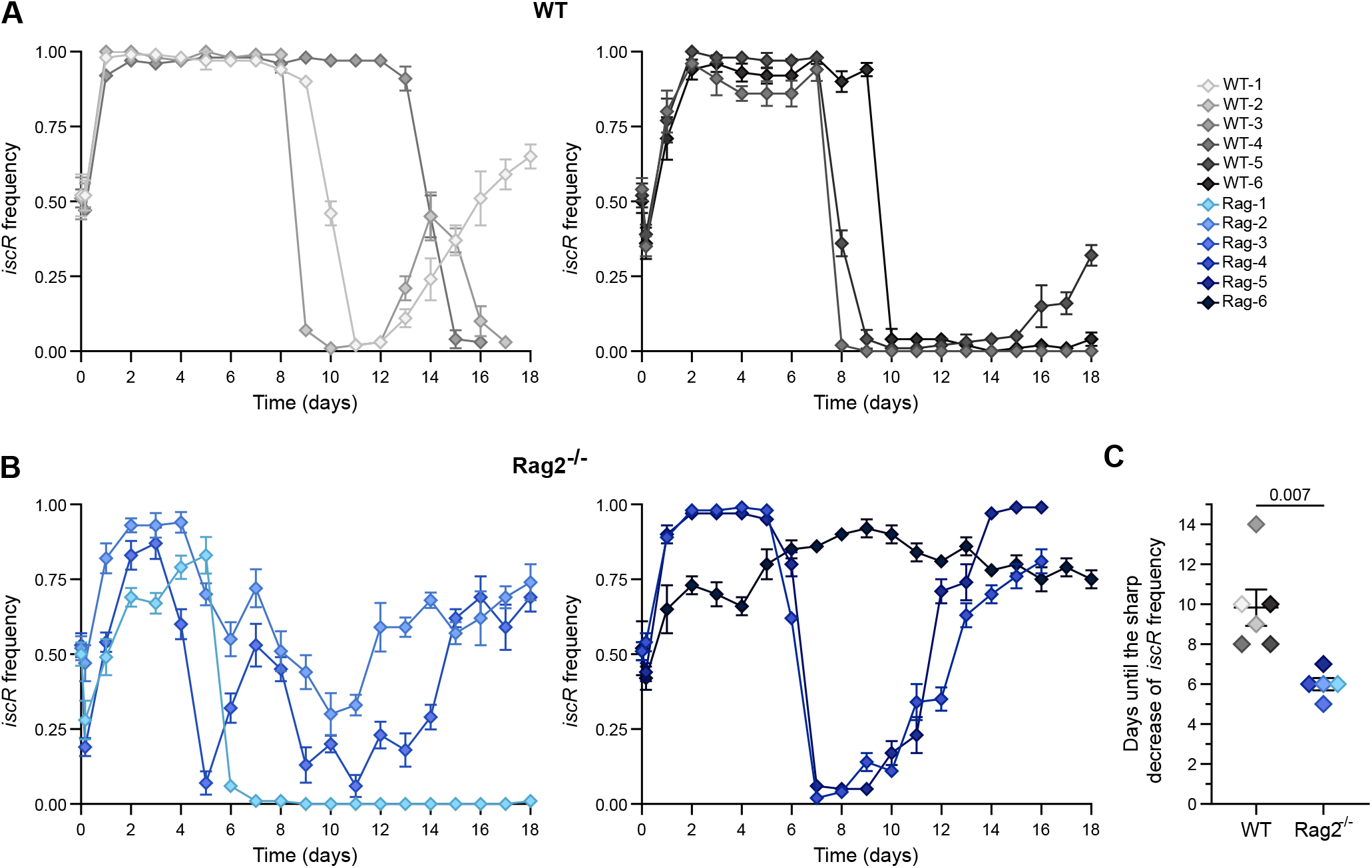
Fluctuating selection of the iron-sulfur cluster regulator *iscR* is modulated by the adaptive immune system. Temporal dynamics of the YFP-labelled streptomycin-resistant *E. coli iscR* when competing against the CFP-labelled streptomycin-resistant wild-type *E. coli* in **(A)** wild-type (WT) (n = 6) and **(B)** *Rag2^-/-^* (n = 6) mice under streptomycin treatment. **(C)** The inversion of selection on *iscR*, measured at the 2 days where a decrease of at least 0.35 in *iscR* frequency was detected, occurs earlier in *Rag2^-/-^* mice (9.8 ± 0.9 in WT, n = 6; and 6.0 ± 0.3 in *Rag2^-/-^* (n = 5); Unpaired t-test with Welch’s correction). In panels A and B the error bars represent the ± 2*SEM, and the left and right plots are biological replicates. In panel C the middle line indicates the mean, and the error bars represent the ± SEM. See also Figure S1 and Table S2.

The timing at which the *iscR* selective change was observed differed between WT and Old mice. This led to the hypothesis that the fluctuating selection could be modulated by the adaptive immune system. A change in selection pressure one-week post-colonization is compatible with a possible response of the adaptive immune system on the colonizing *E. coli*. Indeed, mice can produce IgA specific against *E. coli* (Hapfelmeier et al., 2010), and it is well known that the aging process is accompanied by a decreased effectiveness of the adaptive immune system response (Vaiserman et al., 2017). To test if the fluctuating selection could be modulated by the adaptive immune system, we performed competitions of *iscR* against the wild-type *E. coli* in mice with a compromised immune system. *Rag2^-/-^* mice, which lack mature T and B lymphocytes, are commonly used as a model for immune-compromised hosts (Shinkai et al., 1992; Barroso-Batista et al., 2015), so we tested for fluctuating selection on this mouse model. Again, we observed sharp fluctuations in *iscR* frequency, with a strong selection for *iscR* in the first days of competition followed by an inversion of selection for *iscR* after five to seven days of competition (Figure 2B). When comparing Old, WT, and *Rag2^-/-^* mice, the average selective effect per generation for *iscR* in the first two days did not differ between the groups (Figures S1A and S1B), suggesting that the initial selection for *iscR* is independent of the host physiological state. In contrast, the timing at which the inversion of selection for *iscR* occurred was different between immune-competent and immune-compromised young adult mice (9.8 ± 0.9 SEM vs. 6.0 ± 0.3 SEM, respectively; p = 0.007, Unpaired t-test with Welch’s correction, Figure 2C). Interestingly, no significant differences in total *E. coli* abundance across the different groups of mice were detected, as all groups showed similar numbers of *E. coli* per gram of feces (Figure S1C). This data supports the notion that the adaptive immune system affects the period at which fluctuating selection in the mammalian gut acts on *iscR*.

### *iscR* fluctuating selection is driven by the microbiota and antibiotic treatment

The sharp changes in *iscR* frequency observed over short periods are consistent with a direct change in selection pressure on this gene and unlikely to be driven by the rapid accumulation of mutations in the *E. coli* genomic background. In fact, we have previously shown that the pace of evolution, via the occurrence of metabolic adaptations, is slower in *Rag2^-/-^* mice than in WT mice (Barroso-Batista et al., 2015), making the dramatic recurrent changes in *iscR* frequency observed in *Rag2^-/-^* difficult to reconcile with the rapid emergence of other selected mutations. Such change could be more likely due to microbiota differences, as Old and *Rag2^-/-^* mice have different microbiota compositions than WT mice and these can change rapidly under the antibiotic treatment used (Barroso-Batista et al., 2015; Barreto et al., 2020a). To examine the influence of the microbiota on the competitive fitness of *iscR* we performed competitions in WT and *Rag2^-/-^* germ-free mice which are depleted of microbiota, under the same antibiotic regime used in mice with microbiota. No evidence for fluctuating selection was observed in either WT or *Rag2^-/-^* mice, as in both host genetic backgrounds *iscR* was strongly selected for (Figure 3A). The selective effect per generation estimated from the ratio of abundances of each bacterial clone over time was 0.06 ± 0.01 SEM for WT and 0.05 ± 0.01 SEM for *Rag2^-/-^* mice. The *iscR* clones didn’t show any sharp decrease in frequency in any of the germ-free mice (Figure 3A), suggesting that the presence of a complex microbiota is required for the fluctuating selection of *iscR*.

**Figure 3.**
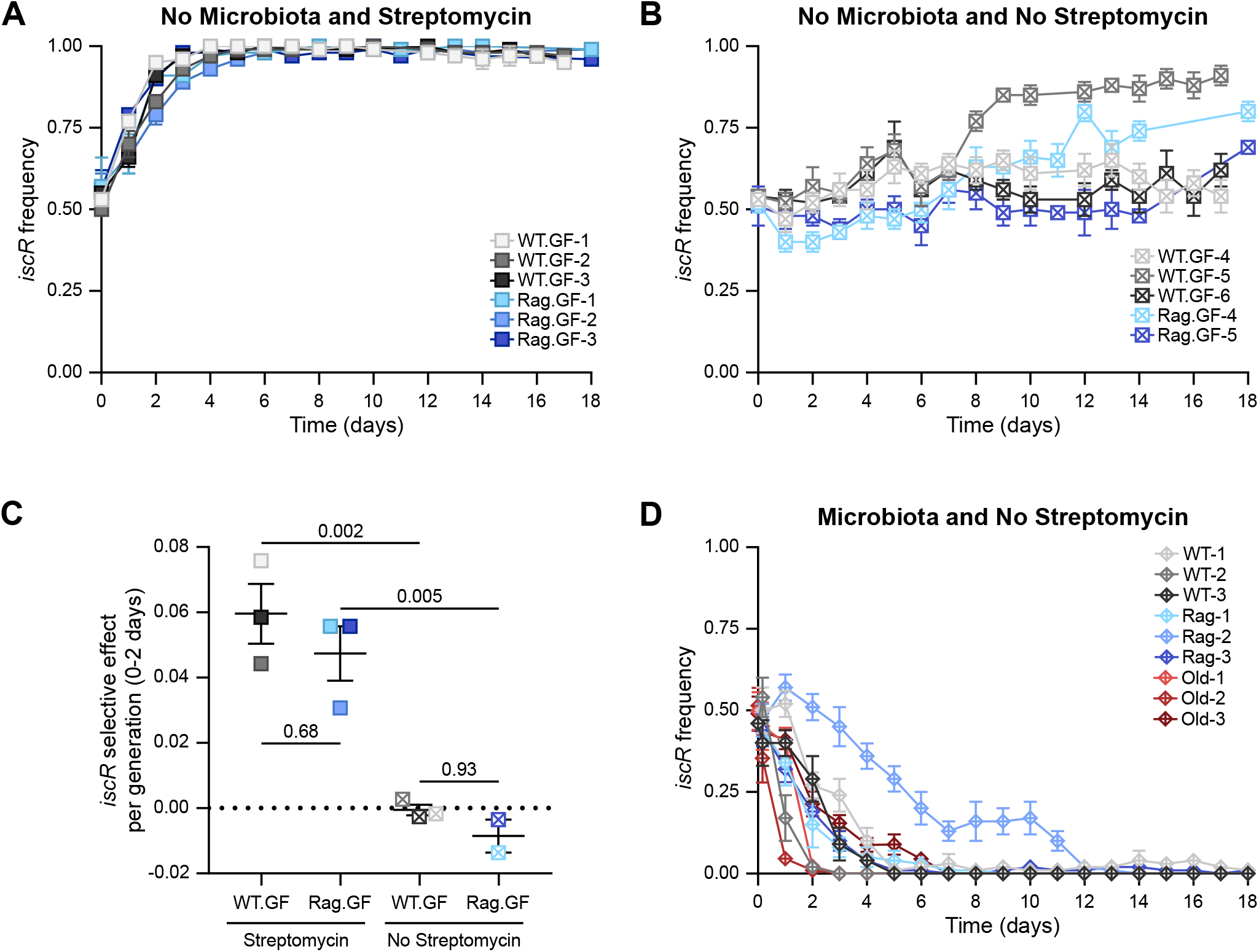
Antibiotic treatment and microbiota are required for fluctuating selection on the iron-sulfur cluster regulator *iscR*. Temporal dynamics of the YFP-labelled streptomycin-resistant *E. coli iscR* when competing against the CFP-labelled streptomycin-resistant wild-type *E. coli* in WT and *Rag2^-/-^* germ-free mice with **(A)** (WT, n = 3; *Rag2^-/-^*, n = 3) and without **(B)** (WT, n = 3; *Rag2^-/-^*, n = 2) streptomycin treatment; **(C)** Selective advantage of *iscR* for the first 2 days of the *in vivo* competitions in WT and *Rag2^-/-^* germ-free mice with and without streptomycin treatment (0.06 ± 0.01 in WT with streptomycin, n = 3; 0.05 ± 0.01 in *Rag2^-/-^* with streptomycin, n = 3; −0.001 ± 0.002 in WT without streptomycin, n = 3; and −0.01 ± 0.01 in *Rag2^-/-^* without streptomycin, n = 2; ANOVA with Šídák’s multiple comparisons test); **(D)** Temporal dynamics of the YFP-labelled *E. coli iscR* when competing against the CFP-labelled wild-type *E. coli* in the gut of WT (n = 3), *Rag2^-/-^* (n = 3), and Old (n = 3) mice without streptomycin treatment. In panels A, B, and D the error bars represent the ± 2*SEM. In panel C the middle line indicates the mean, and the error bars represent the ± SEM. See also Figure S2 and Table S2.

Antibiotics, such as aminoglycosides, are known to affect host physiology independently of the presence of a microbiota (Gopinath et al., 2018). To query if the treatment with the aminoglycoside streptomycin during the *in vivo* competitions in germ-free mice could affect the directional selection observed, we performed competition assays in the presence and absence of streptomycin. No differences in the total loads of *E. coli* were observed in the presence vs. absence of antibiotic treatment (Figure S2A). Surprisingly, no strong signs of directional selection for *iscR* were observed in the absence of antibiotic, as in both WT and *Rag2^-/-^* germ-free mice *iscR* behaved as a neutral mutation during the first week of the competitions (selective effect per generation of - 0.001 ± 0.001 SEM for WT; −0.009 ± 0.005 SEM for *Rag2^-/-^*; Figure 3B). Any signature of fluctuating selection was also absent without streptomycin treatment (Figure 3B). Supporting the observations for a strong selection of *iscR* in the presence of streptomycin, the selective effect of *iscR* in the first two days is significantly higher in the presence of streptomycin than in the absence of streptomycin (Figure 3C). The selective effect is also independent of the mice background, as no differences were observed between WT and *Rag2^-/-^* in both presence or absence of streptomycin (Figures 3C and S2B). These results suggest that the initial strong selection for *iscR* is driven by the antibiotic treatment in both immune-competent and immune-compromised mice lacking microbiota.

Having established that the antibiotic treatment selects for *iscR* in the absence of microbiota, we next tested if the presence of the microbiota but absence of streptomycin could alter the selective effect for *iscR*. Due to colonization resistance, *E. coli* is not able to colonize the mouse gut without a streptomycin pre-treatment, so we performed *in vivo* competitions between *iscR* and wild-type *E. coli* using a modified experimental setup (Leónidas Cardoso et al., 2020) (See also STAR Methods). In this setup, the microbiota is first temporarily perturbed allowing *E. coli* to colonize the mouse gut when streptomycin is withdrawn. In all cohorts of mice tested (Old, WT, and *Rag2^-/-^* mice) we found that *iscR* is strongly selected against in mice with microbiota in the absence of streptomycin (Figure 3D). The estimated selective effect per generation against *iscR* in the first two days of competition was not significantly different in Old, WT, and *Rag2^-/-^* mice, suggesting that *iscR* is equally counter-selected in the absence of antibiotic (−0.09 ± 0.03 SEM for Old; −0.05 ± 0.03 SEM for WT; −0.02 ± 0.02 SEM for *Rag2^-/-^*; Figures S2C and S2D). No differences in the total loads of *E. coli* sampled from the mouse feces were observed between all groups of mice (Figure S2E). These and the previous results show that fluctuating selection on *iscR* is driven by both the microbiota and the streptomycin treatment.

### Iron limitation modulates *iscR* fitness

The fight for iron between host and microbes could be a possible source of fluctuating selection as the host can regulate gut iron concentrations. The host can limit iron availability to prevent pathogenic bacteria from blooming, and iron levels in the colonic region of the mammalian gut can rapidly change (Yilmaz and Li, 2018). Given the importance of IscR in the iron homeostasis of *E. coli*, we tested if the fitness of *iscR* changes under different iron stressing conditions. We compared the growth of *iscR* and wild-type *E. coli in vitro* under iron limitation in the presence or absence of streptomycin. We chose a concentration of streptomycin compatible with the previous measurements in streptomycin-treated mice (Leónidas Cardoso et al., 2020), to mimic this selective pressure occurring in the *in vivo* competitions.

When *iscR* and wild-type *E. coli* were grown in rich Lysogeny Broth (LB) medium, we observed that *iscR* clones have a lower maximum growth rate and carrying capacity than wild-type *E. coli*, irrespective of the presence of streptomycin (maximum growth rate of 0.722 ± 0.005 SEM for *iscR* and 0.79 ± 0.01 SEM for wild-type *E. coli* in the absence of streptomycin, 0.72 ± 0.01 SEM for *iscR* and 0.81 ± 0.01 SEM for wild-type *E. coli* in the presence of streptomycin; carrying capacity of 1.38 ± 0.02 SEM for *iscR* and 1.53 ± 0.01 SEM for wild-type *E. coli* in the absence of streptomycin, 1.43 ± 0.01 SEM for *iscR* and 1.55 ± 0.01 SEM for wild-type *E. coli* in the presence of streptomycin; Figure 4A). We then measured the same fitness traits in iron-limiting conditions, by either adding the host innate immune protein Lipocalin-2, which binds to bacterial siderophores, or by adding an iron chelator (Dipyridyl), which forms a complex directly with iron. We found that, in the presence of Lipocalin-2, *iscR* shows a higher maximum growth rate than wild-type *E. coli*, both in the presence and in the absence of streptomycin: the maximum growth rate estimated was 0.67 ± 0.03 SEM for *iscR* and 0.44 ± 0.02 SEM for wild-type *E. coli* in the absence of streptomycin, 0.65 ± 0.01 SEM for *iscR* and 0.46 ± 0.03 SEM for wild-type *E. coli* in the presence of streptomycin (Figure 4B). Interestingly, the presence of streptomycin also affected the carrying capacity of *iscR*, as *iscR* only achieves higher carrying capacity than the wild-type *E. coli* in the presence of streptomycin: 0.86 ± 0.02 SEM for *iscR* and 0.80 ± 0.03 SEM for wild-type *E. coli* in the absence of streptomycin, 0.92 ± 0.02 SEM for *iscR* and 0.66 ± 0.02 SEM for wild-type *E. coli* in the presence of streptomycin (Figure 4B). Similar to the effect of Lipocalin-2, the presence of the iron chelator Dipyridyl lead to an increased maximum growth rate of *iscR* when compared to the wild-type *E. coli:* 0.36 ± 0.02 SEM for *iscR* and 0.30 ± 0.01 SEM for wild-type *E. coli* in the presence of streptomycin (Figure 4C). In the absence of streptomycin, no significant difference in the maximum growth rate between *iscR* and wild-type *E. coli* was observed (0.28 ± 0.01 SEM for *iscR* and 0.26 ± 0.01 SEM for wild-type). Dipyridyl also caused the carrying capacity of *iscR* to be higher than the wild-type *E. coli*, irrespectively of streptomycin treatment: 0.63 ± 0.02 SEM for *iscR* and 0.37 ± 0.02 SEM for wild-type *E. coli* in the absence of streptomycin, 0.71 ± 0.02 SEM for *iscR* and 0.46 ± 0.02 SEM for wild-type *E. coli* in the presence of streptomycin (Figure 4C). Thus, iron limitation and the antibiotic streptomycin modulate fitness-related traits of *iscR*.

**Figure 4.**
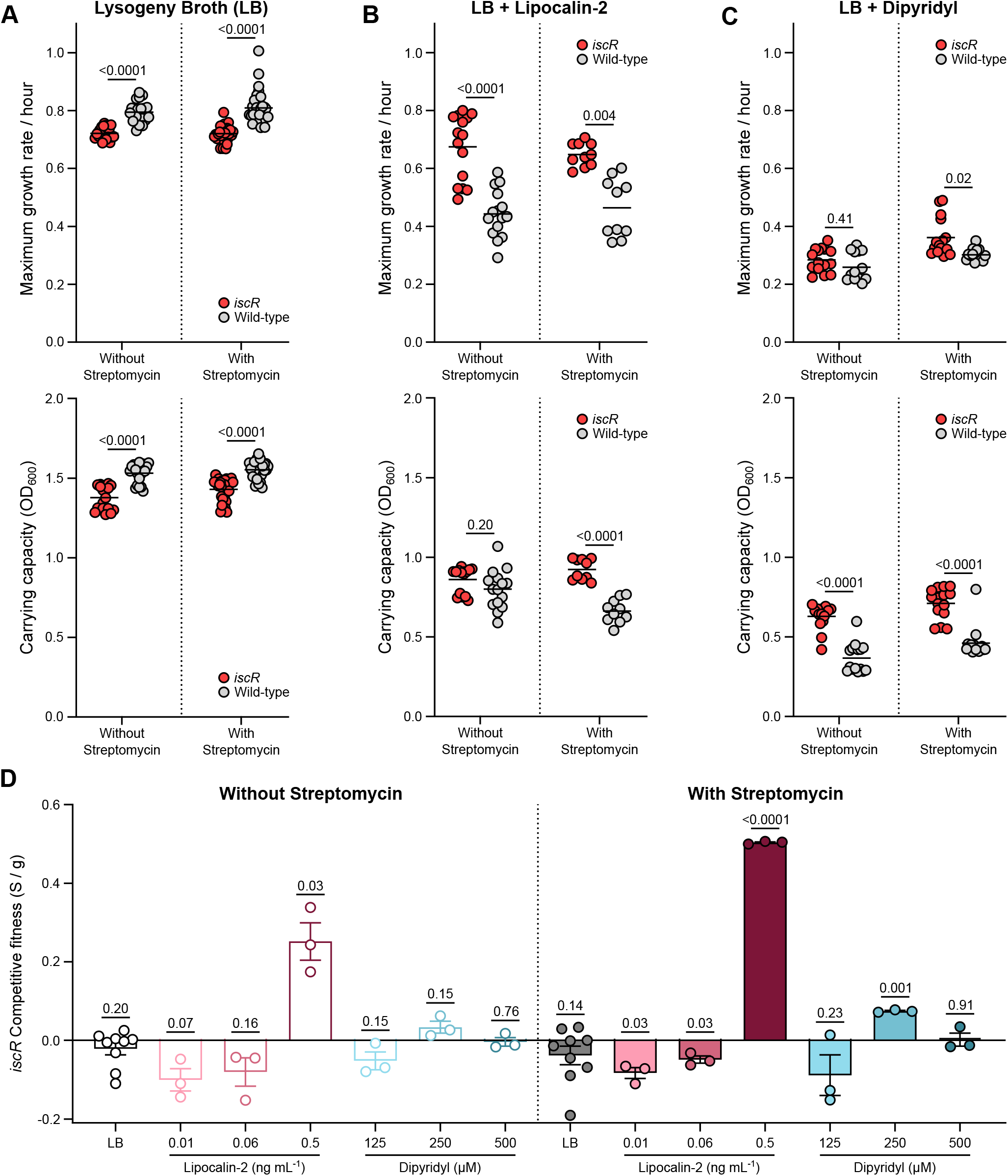
Iron limitation caused by the host protein Lipocalin-2 or the iron chelator Dipyridyl modulates growth traits and competitive fitness of the iron-sulfur regulator *iscR*. **(A)** Maximum growth rate per hour (top panel) and carrying capacity (bottom panel) of *iscR* and wild-type *E. coli* in LB with (n = 30 for both *iscR* and wild-type) and without (n = 20 for both *iscR* and wild-type) 16 μg ml^-1^ streptomycin; **(B)** Maximum growth rate per hour (top panel) and carrying capacity (bottom panel) of *iscR* and wild-type *E. coli* in LB supplemented with 0.5 ng ml^-1^ Lipocalin-2 (n = 10 for both *iscR* and wild-type with streptomycin; n = 15 for both *iscR* and wild-type without streptomycin); **(C)** Maximum growth rate per hour (top panel) and carrying capacity (bottom panel) of *iscR* and wild-type *E. coli* in LB supplemented with 500 μM Dipyridyl (n = 15 for both *iscR* and wild-type with streptomycin; n = 15 for both *iscR* and wild-type without streptomycin); **(D)** Competitive index per generation of *iscR* against wild-type *E. coli* in LB in the absence (n = 9) or presence (n = 9) of 16 μg ml^-1^ streptomycin with varying Lipocalin-2 (n = 3 in all concentrations tested) and Dipyridyl (n = 3 in all concentrations tested) concentrations. In panels A, B, and C, the line represents the mean, and a Kruskal-Wallis test with Dunn’s multiple comparison test was used. In panel D the middle line indicates the mean, the error bars represent the ± SEM, and a One sample t-test with a theoretical mean of 0 was used. See also Table S3 and S4.

We next tested how these stresses would influence the competitive fitness of *iscR*. Since iron can be a public good (Griffin et al., 2004), the effect of the *iscR* allele could be different when the clones grow alone vs in competition (Özkaya et al., 2018). *In vitro* competitions of *iscR* against the wild-type *E. coli* showed that while the mutation was neutral in the presence or absence of streptomycin in LB (Figure 4D), it becomes beneficial under iron limitation. This competitive fitness benefit of *iscR* was dependent on the concentration of Lipocalin-2 or Dipyridyl and occurs in the presence or absence of streptomycin (Figure 4D). Remarkably, at high concentrations of Lipocalin-2, with streptomycin, the advantage of *iscR* was very large (0.504 ± 0.002 SEM for 0.5 ng mL^-1^ of Lipocalin-2), but at lower concentrations of Lipocalin-2 the mutation was deleterious (−0.08 ± 0.01 SEM for 0.01 ng mL^-1^ of Lipocalin-2) (Figure 4D). This change in the sign of the selection coefficient predicts that the mutation should undergo fluctuating selection in an environment where the antibiotic is present and the Lipocalin-2 levels would change. The interaction between iron limitation and the antibiotic on the fitness of *iscR* was also seen in medium supplemented with Dipyridyl (Figure 4D).

These results show that under iron limiting environments the fitness of *iscR* can substantially change and its competitive ability can be modulated either positively or negatively in the presence of streptomycin.

### Levels of lipocalin-2 in the mouse gut correlate with the abundance of *iscR*

The predictions of the *in vitro* fitness measures on *iscR* led us to search for an association between the levels of Lipocalin-2 in feces and the abundance of *iscR* in the mouse gut. If the concentration of Lipocalin-2 changes in the mouse gut, then such changes could potentially contribute to the fluctuating selection dynamics of *iscR* observed *in vivo*. To test this hypothesis, we measured the concentration of Lipocalin-2 in feces of all mice used in the *in vivo* competitions. We find that the time series of fecal Lipocalin-2 levels in all mice show substantial temporal fluctuations that can span 3 Logs (Figure 5A and S3). As expected, in Old mice the average level of fecal Lipocalin-2 is higher than in the other mouse cohorts, but occasionally high levels of Lipocalin-2 were observed in WT and *Rag2^-/-^* mice (Figure S3). We also found substantial fluctuations in this host trait in ex-germ-free mice that were mono-colonized with *E. coli*, irrespective of antibiotic treatment (Figure S4). Streptomycin treatment did not cause any significant increase in the mean Lipocalin-2 concentrations (Figure S5A). Thus, rapid changes of fecal Lipocalin-2 can be experienced in the mouse gut with or without antibiotic treatment. Fluctuations in fecal Lipocalin-2 concentration can also be detected in unmanipulated mice, irrespective of the host age or genetic background (Figure S5B). The data from all time points across all mice also indicates that the microbiota and antibiotic could modulate the temporal mean levels of Lipocalin-2 in the mouse gut. These results therefore suggest that sufficient temporal variation in the levels of fecal Lipocalin-2 exists which could contribute to changes in the dynamics of *iscR in vivo*.

**Figure 5.**
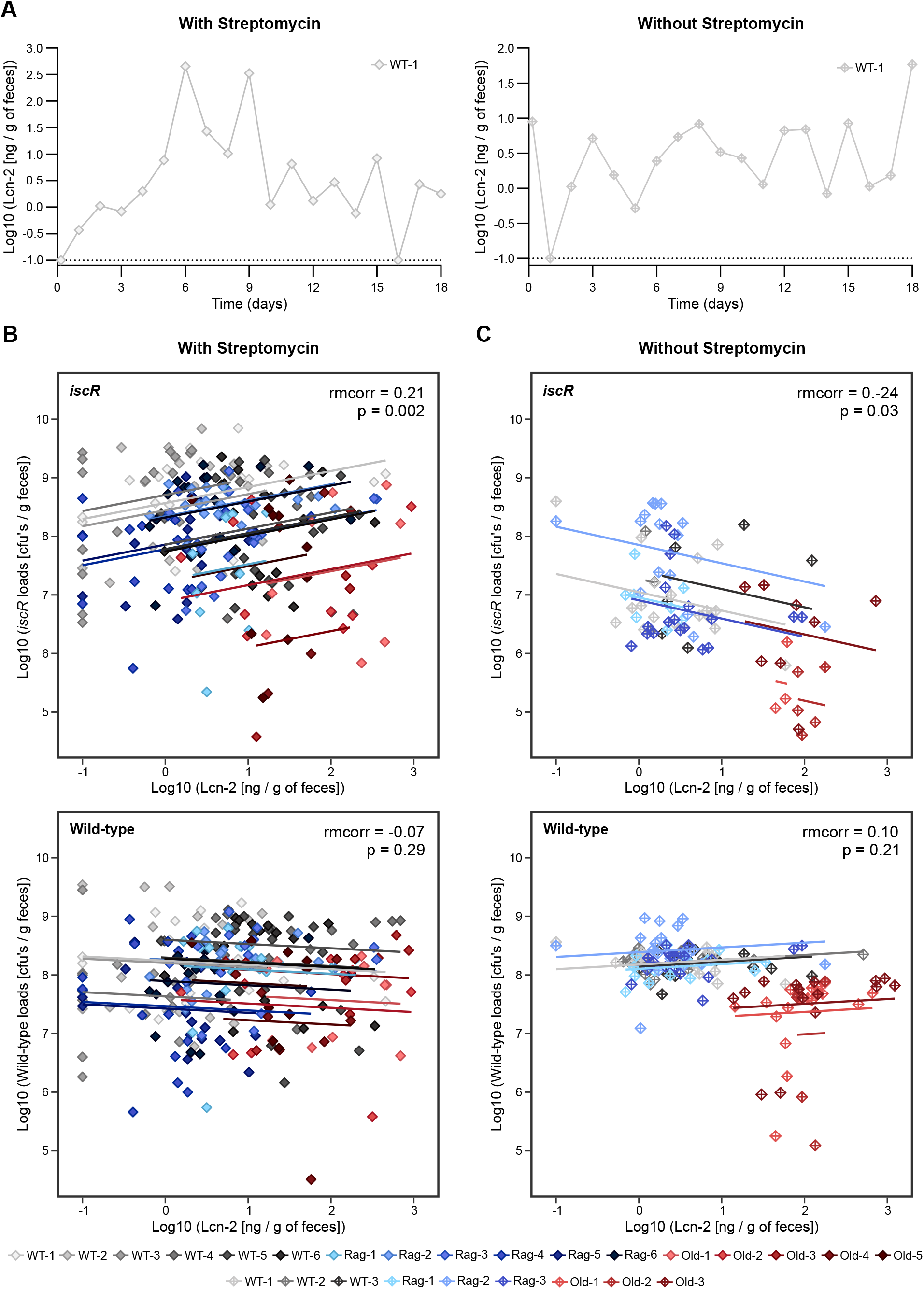
The iron-sulfur cluster *iscR* abundances correlate with fecal Lipocalin-2 concentrations in an antibiotic-dependent manner. **(A)** Representative mice showing the fecal Lipocalin-2 levels throughout time in the presence (left panel) and absence (right panel) of streptomycin treatment; **(B)** Repeated measures correlation between *iscR* (top panel) or wild-type *E. coli* (bottom panel) abundance and fecal Lipocalin-2 levels in mice with microbiota treated with streptomycin (n = 17); **(C)** Repeated measures correlation between *iscR* (top panel) or wild-type *E. coli* (bottom panel) abundance and fecal Lipocalin-2 levels in mice with microbiota not treated with streptomycin (n = 9). See also Figures S3, S4, S5, and Table S2.

We tested for correlations between the levels of fecal Lipocalin-2 and the abundance of either *iscR* or wild-type *E. coli* during the *in vivo* competition experiments. A repeated measures correlation (Bakdash and Marusich, 2017) using the measured samples from animals with or without streptomycin treatment, showed that in the presence of streptomycin there was a significant positive correlation between *iscR* loads and fecal Lipocalin-2 (rmcorr = 0.21, p = 0.002; Figure 5B), suggesting that, in this condition, the fitness of *iscR* increases when the levels of lipocalin-2 increase. This result was specific to *iscR* as no correlation was detected between the loads of wild-type *E. coli* and fecal Lipocalin-2 (Figure 5B). Remarkably, in mice without antibiotic treatment, the abundances of *iscR* were significantly negatively correlated with fecal Lipocalin-2 (rmcorr = −0.24, p = 0.03; Figure 5C), and again no correlation between wild-type *E. coli* loads and fecal Lipocalin-2 was detected (Figure 5C). These results are consistent with the expectations from the *in vitro* experiments for fitness traits and competitive fitness, where the advantage or disadvantage of *iscR* was dependent on the concentration of Lipocalin-2 and modulated by streptomycin. However, the *in vitro* experiments lack a complex microbiota, therefore we next performed the same analysis on the data obtained in germ-free mice. No correlations between either *iscR* or wild-type loads and Lipocalin-2 were detected, either in the ex-germfree mice with and without streptomycin treatment (Figures S5C and S5D). This result shows that the microbiota is necessary to modulate the change in fitness of the *iscR* mutant with the fecal Lipocalin-2 levels in the gut and that the antibiotic acts on the ecosystem to alter such association.

## DISCUSSION

The availability of iron in the mammalian gut is of utmost importance for either the host or the microbes that inhabit this dynamic environment (Yilmaz and Li, 2018). It is well known that both abiotic (*e.g*. antibiotic treatment) and biotic (*e.g*. chronic inflammation) factors can perturb iron availability in the mammalian gut environment, with consequences for the host and their resident bacteria (Chassaing et al., 2012; Yilmaz and Li, 2018; Lu et al., 2019; Ng et al., 2019). Therefore, in the rapidly changing environment of the mammalian gut microbiota, where iron availability can change very rapidly, bacteria need to quickly alter their regulatory circuits in response to environmental cues. Recently, we described the emergence of mutations in the iron-cluster regulator IscR in aged mice (Barreto et al., 2020a), more specifically in the cysteine residues that are essential for the assembly of IscR iron-sulfur cluster. Given *iscR* major role in iron homeostasis (Santos et al., 2015; Esquilin-Lebron et al., 2021) we tested the competitive ability of iscR mutants in the gut commensal bacterium *E. coli* in mice of different ages and immunocompetence.

Understanding how bacterial evolution is affected by fluctuating environments has been intensively studied. However, most studies focused on artificially fluctuating environments (Tagkopoulos et al., 2008; Beaumont et al., 2009; Sandberg et al., 2017; Turner et al., 2020; Nguyen et al., 2021) or theoretical predictions (Melbinger and Vergassola, 2015; Sæther and Engen, 2015; Novak and Barton, 2017; Erickson et al., 2017). Here, using a natural environment, we show that a C92S mutation in *iscR* is under fluctuating selection in the mammalian gut. Given its role as a sensor of environmental conditions that can affect the cellular iron-sulfur cluster pool, IscR has been implicated in *E. coli* adaptation to fluctuating environments (Esquilin-Lebron et al., 2021). A clear example of such conditions is its role in maintaining cell-to-cell variability during anaerobic to aerobic switch, enhancing *E. coli* fitness (Carey et al., 2018). In the gut the observed frequency dynamics of *iscR* are consistent with fluctuating selection acting on the C92S mutation in WT, *Rag2^-/-^* and aging mice (Figures 1 and 2). Furthermore, our findings show that the adaptive immune system can influence the period of fluctuating selection (Figure 2C). The relationship between the adaptive immune system and iron homeostasis has been studied, with dysregulation of iron homeostasis affecting both T and B lymphocytes (Mu et al., 2021). It is still however unclear if immunodeficiency, such as the lack of T and B lymphocytes in *Rag2^-/-^* mice, can affect host iron homeostasis. With aging the development of immunosenescence is well known (Vaiserman et al., 2017), together with chronic inflammation that can lead to the development of anemia of chronic inflammation (Wessling-Resnick, 2010), which disrupts host iron homeostasis. Given the relevance of IscR in the iron circuitry of *E. coli*, our results support the notion that the adaptive immune system is an important factor affecting the action of natural selection acting on this trait in the gut.

The gut microbial community is also known to affect iron homeostasis, as the absence of microbes can lead to depletion of iron (Reddy et al., 1972; Deschemin et al., 2016). Our results in ex-germ-free mice are partially consistent with this, as a strong beneficial effect of the mutation in *iscR* was seen but only in the presence of streptomycin (Figure 3A). These results were observed in both WT and *Rag2^-/-^* ex-germ-free mice, suggesting that the combination of a gut microbial community and antibiotic treatment dictates how natural selection acts on *iscR* in immuno-competent and immuno-compromised hosts. Supporting this idea, our results in mice with a microbiota but no antibiotics revealed that *iscR* is continuously selected against, with no signs of fluctuating selection irrespectively of mice age or immune competence (Figure 3D). These results may explain why the emergence of mutations in the cysteine residues of IscR was only detected during short-term evolution of *E. coli* in the presence (Barreto et al., 2020a) but not absence of antibiotic treatment (Barreto et al., 2020b) in aging hosts.

Antibiotic tolerance has also been related with the regulation of iron-sulfur clusters. IscR regulates the expression of the *isc* operon (Schwartz et al., 2001), composed by *iscRSUA-hscBA-fdx*. The *isc* operon is repressed when IscR assembles its iron-sulfur cluster. Accordingly, deletion of *iscR* leads to increased expression of *iscSUA* (Schwartz et al., 2001), and IscR mutants that are unable to assemble its iron-sulfur cluster, such as the mutation C92S studied here, show no repression of the *isc* operon (Giel et al., 2013). Interestingly, overexpression of *iscU* confers an advantage to *E. coli* in the presence of the aminoglycoside streptomycin, as it decreases protein aggregation induced by streptomycin treatment (Ling et al., 2012). In addition, an *iscUA* mutant was also shown to be resistant to the aminoglycoside gentamycin (Ezraty et al., 2013). Although we use *E. coli* strains that are resistant to streptomycin, they are still able to uptake this antibiotic, which could in part explain the *in vivo* competitive advantage seen for *iscR* when competing against the wild-type *E. coli* in the presence of streptomycin. However, the *in vitro* results showing no advantage for *iscR* in the presence of streptomycin alone (Figure 4) suggest that it is not a direct interaction between *iscR* and streptomycin that drives the advantage of *iscR in vivo*.

Using an iron limiting environment, either by the addition of the host innate immune protein Lipocalin-2 or the iron chelator Dipyridyl, we show that the presence of streptomycin in the culture media can modulate both the growth traits and competitive fitness of *iscR* (Figure 4). Aminoglycosides, such as streptomycin, can form a complex with iron that can lead to ROS formation and consequently oxidative stress (Ezraty and Barras, 2016). In addition, the *isc* operon is activated under iron limiting and oxidative stress conditions (Yeo et al., 2006), when IscR is predominantly in its clusterless form. Given that the *iscR* mutant used in this work is locked in its clusterless form, the modulation of growth traits and competitive fitness of *iscR* could be a consequence of an additive effect from the two processes described above. We also observed the interaction between Lipocalin-2 and streptomycin *in vivo*, as the correlation between *iscR* abundance and fecal Lipocalin-2 can change from positive to negative depending on the antibiotic treatment (Figure 5B), in agreement with the effects measured *in vitro*, highlighting the key role of Lipocalin-2 and streptomycin for *iscR* fluctuating selection dynamics. Our data does not however exclude that other factors, either host-related or microbe-related, may also be important for the fluctuating selection acting on *iscR*.

It has been described that intestinal Lipocalin-2 expression is coordinated by the circadian cycle and by the attachment in the intestinal surface of segmented filamentous bacteria (SFB), a member of the mouse gut microbiota (Brooks et al., 2021). In addition, in germ-free mice, Lipocalin-2 shows no rhythmic expression, and in the presence of the aminoglycoside streptomycin the colonization with SFB is reduced leading to a likely abolishment of Lipocalin-2 rhythmic expression (Brooks et al., 2021). In contrast, our daily measurements of fecal Lipocalin-2 in mice with a complex microbiota show substantial temporal fluctuations that sometimes can span 3 Logs (Figures 5A, S3, and S5A), irrespectively of the antibiotic treatment. We also observed no differences in the fecal Lipocalin-2 levels for ex-germ-free mice with or without antibiotic treatment (Figure S4 and S5A). Despite the similar temporal fluctuations of Lipocalin-2 in the mice feces, it is likely that the presence of a microbiota may alter the levels of Lipocalin-2 inside the mouse intestine where the mutant *iscR* and wild-type *E. coli* compete (Cohen and Conway, 2015). It is conceivable that a decrease in iron availability at those sites, could lead to the fluctuating selection dynamics observed in mice with microbiota.

In conclusion, to the best of our knowledge, our work is the first to detect a mode of natural selection – fluctuating selection – so far not described to shape bacteria evolution in the mammalian gut. The data highlights how subtle the interactions between the host immune system, the microbiota community, and antibiotic treatment can be in tempering the way natural selection acts on critical bacterial processes, such as iron homeostasis. As the mammalian gut is known to be a dynamic environment, it is likely that during bacterial evolution other mutations that target important bacterial processes may undergo fluctuating selection. Thus, understanding the extent to which fluctuating selection shapes within-species diversity in the microbiota ecosystem is an important direction for future work. The increasing number of longitudinal studies focusing on bacteria evolution in the mammalian gut should aid in accomplishing such a task.

## ACKNOWLEDGMENTS

We thank Gordo’s lab members for the helpful discussions throughout the work and for critically reading this manuscript. We thank Nelson Frazão, Ricardo Ramiro and Paulo Durão for the help with *in vivo* competitions. This work was supported by Global Grant for Gut Health-623877 to I.G., and partially supported by ONEIDA project (LISBOA-01-0145-FEDER-016417) co-funded by FEEI - “Fundos Europeus Estruturais e de Investimento” from “Programa Operacional Regional Lisboa 2020” to I.G.. H.C.B. was the recipient of a doctoral fellowship (PD/BD/128429/2017) from the FCT (“Fundação para a Ciência e a Tecnologia”). This work was developed with the support from the research infrastructure Congento, co-financed by Lisboa Regional Operational Programme (Lisboa2020), under the PORTUGAL 2020 Partnership Agreement, through the European Regional Development Fund (ERDF) and Fundação para a Ciência e Tecnologia (Portugal) under the project LISBOA-01-0145-FEDER-022170. The funders had no role in study design, data collection and analysis, decision to publish, or preparation of the manuscript.

## AUTHOR CONTRIBUTIONS

Conceptualization, H.C.B. and I.G.; Methodology, H.C.B. and I.G.; Validation, H.C.B. and I.G.; Formal Analysis, H.C.B. and I.G.; Investigation, H.C.B. and B.A.; Resources, I.G.; Writing – Original Draft, H.C.B. and I.G.; Writing – Review & Editing, H.C.B. and I.G.; Visualization, H.C.B. and I.G.; Funding Acquisition and Supervision, I.G.

## STAR METHODS

### LEAD CONTACT AND MATERIALS AVAILABILITY

Further information and requests for resources and reagents should be directed to and will be fulfilled by the Lead Contact, Isabel Gordo.

### MATERIALS AVAILABILITY

This study did not generate new unique reagents.

### DATA AND CODE AVAILABILITY

Genome sequencing data have been deposited with links to BioProject accession number PRJNA794030 in the NCBI BioProject database (https://www.ncbi.nlm.nih.gov/bioproject/). Any additional information required to reanalyze the data reported in this paper is available from the lead contact upon request. This paper does not report original code.

### EXPERIMENTAL MODEL AND SUBJECT DETAILS

#### Mice

This research project was ethically reviewed and approved by both the Ethics Committee and the Animal Welfare Body of the IGC (license reference: A009.2018), and by the Portuguese National Entity that regulates the use of laboratory animals (DGAV – Direção Geral de Alimentação e Veterinária (license reference: 009676). All experiments conducted on animals followed the Portuguese (Decreto-Lei n° 113/2013) and European (Directive 2010/63/EU) legislations, concerning housing, husbandry, and animal welfare. For the *in vivo* competitions in the presence of microbiota, female C57BL6/6J (2-3 months of age for WT mice and 19-30 months of age for Old mice) and C57BL6/6J *Rag2^-/-^* (2-3 months of age) specific pathogen-free mice were used. These animals were bred and maintained at the Instituto Gulbenkian de Ciência rodent facility. For the *in vivo* competitions in the absence of microbiota, female and male germ-free C57BL6/6J (5-6 months of age) and C57BL6/6J *Rag2^-/-^* (5-6 months of age) mice were used. These animals were bred and raised at the gnotobiology unit, in axenic isolators (La Calhene/ORM) and later transferred into sterile ISOcages (Tecniplast). All animals used in this work were single housed with access to food and water *ad libitum*.

#### Bacteria

All bacterial strains used in this work were derived from *E. coli* K-12 MG1655, strains DM08-YFP and DM09-CFP as previously described (Barreto et al., 2020a). Construction of the wild-type strain with a gat-negative background was achieved through the selection of spontaneous gat-negative mutants by incorporating 10 mM D-Arabitol in the selection plates. The *iscR* mutant was previously isolated (Barreto et al., 2020a), carrying an IS insertion in the galactitol operon and a non-synonymous substitution in *iscR*, with the cysteine in position 92 replaced by a serine. The wild-type *E. coli* carries an IS insertion in the galactitol operon. Both strains are resistant to streptomycin and ampicillin and their genome was sequenced at the IGC Genomics facility using an Illumina NextSeq 500. Sequencing adapters were autodetected and removed using *fastp* (Chen et al., 2018). Raw reads were trimmed from both sides, using window sizes of 4 base pairs across which the average base quality had to reach a minimum value of 20 to be retained. Trimmed reads were retained if they reached a minimum length of 100 bps and consisted of at least 50% base pairs that had phred scores of at least 20. Mutations were then identified using the BRESEQ pipeline (Deatherage and Barrick, 2014) and each strain carries only the mutations described above. *E. coli* was routinely cultured in Lysogeny-Broth (LB) at 37°C with aeration except when otherwise indicated.

### METHOD DETAILS

#### Conservation analysis of IscR residues

Conservation analysis of IscR residues was performed using the Consensus Finder web tool (http://kazlab.umn.edu/) (Jones et al., 2020). As an input, we used the amino acid sequence of IscR obtained from UniProt (Accession code: P0AGK8). Consensus Finder default options were used, except for the maximum sequences for BLAST search (set to 10000), the maximum e-value for BLAST search (set to 10^-30^), and CD-Hit redundancy (set to 1). From this analysis, we obtained a total of 8284 hits, and the conservation frequency for each amino acid can be found in Table S1.

#### Competitive fitness assay *in vivo*

To assess the *in vivo* dynamics of YFP-labeled *iscR*, we performed *in vivo* competition assays in a gat-negative background against the CFP-labeled wild-type at a ratio of 1 to 1. For the competitive fitness assays in the presence of streptomycin, the mice were given autoclaved water containing streptomycin (5 g L^-1^) for one day. After 4 hours of starvation for water and food, the animals were orally gavaged with 100 μL of a 10^8^ suspension of the 1 to 1 *E. coli* mixture. After gavage, both water containing streptomycin (5 g L^-1^) and food were made accessible to the animals. During the course of the experiment, the water always contained streptomycin (5 g L^-1^) and was changed every 3 days. For the competitive fitness assays in the absence of streptomycin and absence of microbiota, the procedure was the same as described above but without adding streptomycin to the autoclaved water. For the competitive fitness assays in the absence of streptomycin and the presence of microbiota, the mice were given autoclaved water containing streptomycin (5 g L^-1^) for seven days, followed by two days of regular autoclaved water (Leónidas Cardoso et al., 2020). The mice were then starved for food and water for 4 hours and orally gavaged with 100 μL of a 10^8^ suspension of the 1 to 1 *E. coli* mixture. After gavage, both food and regular autoclaved water were made accessible to the animals. In all competitive fitness assays, fecal pellets were collected daily, diluted in PBS 1X, and plated in LB agar supplemented with streptomycin (100□μg□mL^-1^) to select for the *E. coli* strains or stored with 15 % glycerol at −80°C for future analysis. After incubating the plates overnight at 37°C in aerobiosis, the total loads and the frequency of YFP and CFP bacteria was assessed using a fluorescent stereoscope (Zeiss Stereo Lumar V12). The selection coefficient of *iscR in vivo* was estimated as the slope of the linear regression of ln(freq_*iscR*_/freq_wild-type_) for 2 days after the beginning of the competition.

#### Measurement of *E. coli* fitness traits

*iscR* and wild-type *E. coli* were streaked from the frozen stock into LB agar supplemented with 100 μg ml^-1^ streptomycin and incubated at 37 °C for 24 hours. A minimum of 5 individual colonies were selected from each strain and grown overnight at 37 °C without agitation in LB supplemented with 100 μg ml^-1^ streptomycin. Appropriate dilutions were then performed to reach an OD_600_ of 0.05, followed by the addition in a 96-well plate of 10 μL of culture into 200 μL of LB supplemented with different combinations of streptomycin, Lipocalin-2, and Dipyridyl, as indicated in the legend of Figure 4. The growth dynamics were then obtained by measuring OD_600_ every 20 minutes for 24 hours in a Synergy H1 microplate reader (BioTek) at 37 °C with agitation. The maximum growth rate per hour was obtained using the R package *growthrates* version 0.8.2 (Hall et al., 2014) with a minimum of 4 time points, a smoothness of 0.5, and a quota of 0.95. The carrying capacity was determined by the maximum OD_600_ during the 24 hours growth.

#### Competitive fitness assay *in vitro*

*iscR* and wild-type *E. coli* were streaked from the frozen stock into LB agar supplemented with 100 μg ml^-1^ streptomycin and incubated at 37 °C for 24 hours. A minimum of 3 individual colonies was selected from each strain and grown overnight at 37 °C without agitation in LB supplemented with 100 μg ml^-1^ streptomycin. Appropriate dilutions were then performed to reach an OD_600_ of 0.05, and both strains were mixed at a ratio of 1:1, followed by the addition in a 96-well plate of 10 μL of the mixture into 200 μL of LB supplemented with different combinations of streptomycin, Lipocalin-2, and Dipyridyl, as indicated in the legend of Figure 4. The 96-well plate was then incubated at 37°C with agitation for 24 hours. To determine the initial and final counts of *iscR* and wild-type *E. coli*, the initial mixture and the culture after 24 hours of growth were properly diluted and plated in LB agar plates supplemented with 100 μg ml^-1^ Streptomycin. After incubating the plates overnight at 37°C in aerobiosis, the frequency of YFP and CFP bacteria was assessed using a fluorescent stereoscope (Zeiss Stereo Lumar V12). The selection coefficient of each competition was estimated as the per generation difference in the ratio of the *iscR* and the wild-type *E. coli* after 24□hours: s□=□ln[Rf/Ri]/t, where t is the number of generations and Rf and Ri are the final and initial ratios between *iscR* and wild-type *E. coli*, respectively.

#### Measurement of fecal Lipocalin-2

To measure Lipocalin-2 in the feces, we used the fecal samples obtained in the *in vivo* competition experiments that were frozen at −80 °C. We then scraped the homogenized fecal pellet, vortex for 5 minutes, and centrifuge for 15 minutes at 18000 g, 4°C. From the recovered supernatant we assayed the concentration of lipocalin-2 by ELISA as indicated by the manufacturer (Mouse lipocalin-2/NGAL DuoSet ELISA, R&D Systems).

#### Repeated measures correlation

To test for an association between the levels of fecal Lipocalin-2 and the loads of *E. coli* in the gut we performed a repeated measures correlation using the R package *rmcorr* (Bakdash and Marusich, 2017). This method accounts for the paired repeated measures and variation between individuals. In this analysis, we used the sampling points from the *in vivo* competitive fitness experiments, in which the participant was each mouse and the two measures were the fecal Lipocalin-2 and the loads of *iscR* or the loads of the wild-type *E. coli*. For this analysis, the default parameters were used.

### QUANTIFICATION AND STATISTICAL ANALYSIS

All statistical analyses were conducted in GraphPad Prism or the statistical software R (R Core Team, 2018). Detailed statistics for all the experiments can be found in the figure legends and/or in the manuscript together with the n and definitions of center and dispersion. In all figures, n represents the number of animals that were used, except for Figure 4 where n represents the number of biological independent assays. In all figures, the mean and standard error of the mean (SEM) were used, except otherwise indicated in the manuscript and/or figure legends. The normality of the groups was checked using the Shapiro-Wilk normality test. Statistical significance was defined for p < 0.05 in all comparisons and calculated as described in the manuscript and/or figure legends.

**Table S1. Conservation frequency of IscR residues. Related to Figure 1.** In bold are indicated the amino acid residues with higher conservation for that position. The highlighted positions indicate the amino acids essential for the assembly of the iron-sulfur cluster.

**Table S2. *iscR* frequency, fecal Lipocalin-2 concentration and abundance of total *E. coli, iscR*, and wild-type *E. coli* in the *in vivo* competitions. Related to Figure 1, 2, 3, 5, S1, S2, S3, S4, and S5.** NS, no sample; BDL, below detection limit; SEM, Standard Error of the Mean.

**Table S3. Maximum growth rate per hour and carrying capacity of *iscR* and wild-type *E. coli in vitro*. Related to Figure 4.**

**Table S4. Competitive index per generation of *iscR* against wild-type *E. coli in vitro*. Related to Figure 4.**

