## Supplementary Figures for "Gut mélange à trois: fluctuating selection modulated by microbiota, host immune system, and antibiotics"

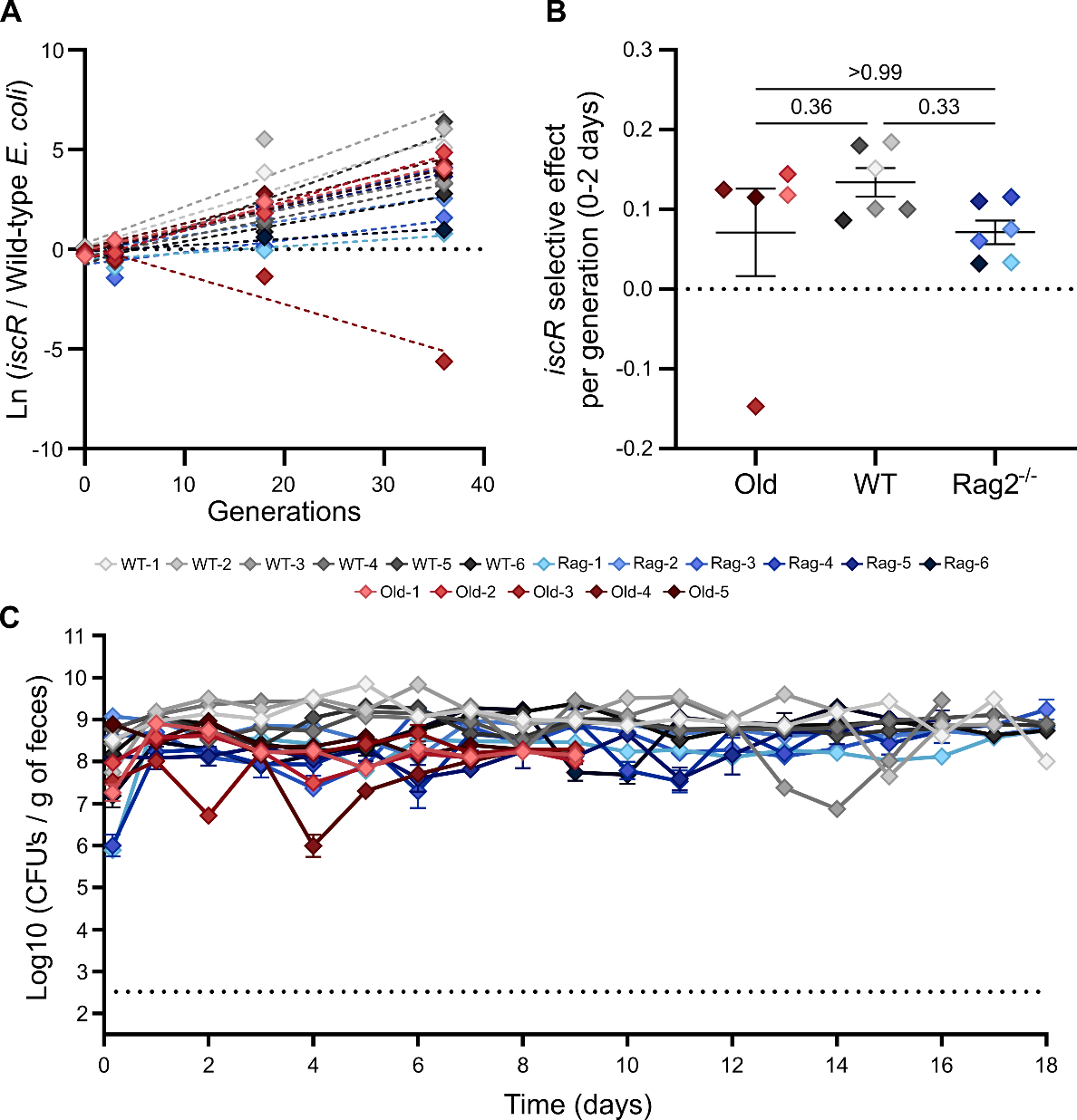

**Figure S1. The selective effect of *iscR* and *E. coli* total loads are independent of the host physiological state*.* Related to Figure 2. (A)** The selective effect of *iscR* after two days of competition in Old (n = 5), WT (n =6), and *Rag2^-/-^* (n = 6) mice is inferred from the slope of the linear regression, corresponding to the selection coefficient. The dashed lines represent the best-fit linear regression; **(B)** Selective effect of *iscR* after two days of competition in Old (0.07 ± 0.05, n = 5), WT (0.13 ± 0.02, n = 6) and *Rag2^-/-^* (0.07 ± 0.01, n = 6) mice. The middle line indicates the mean, the error bars represent ± SEM. For statistical significance the ANOVA with Tukey’s multiple comparison test was used; **(C)** *E. coli* loads per gram of feces throughout the competitions of *iscR* against the wild-type *E. coli* in Old (n = 5), WT (n = 6), and *Rag2^-/-^* (n = 6). Error bars represent the ± 2*SEM. In panel C, the dashed lines indicate the limit of detection. See also Table S1.

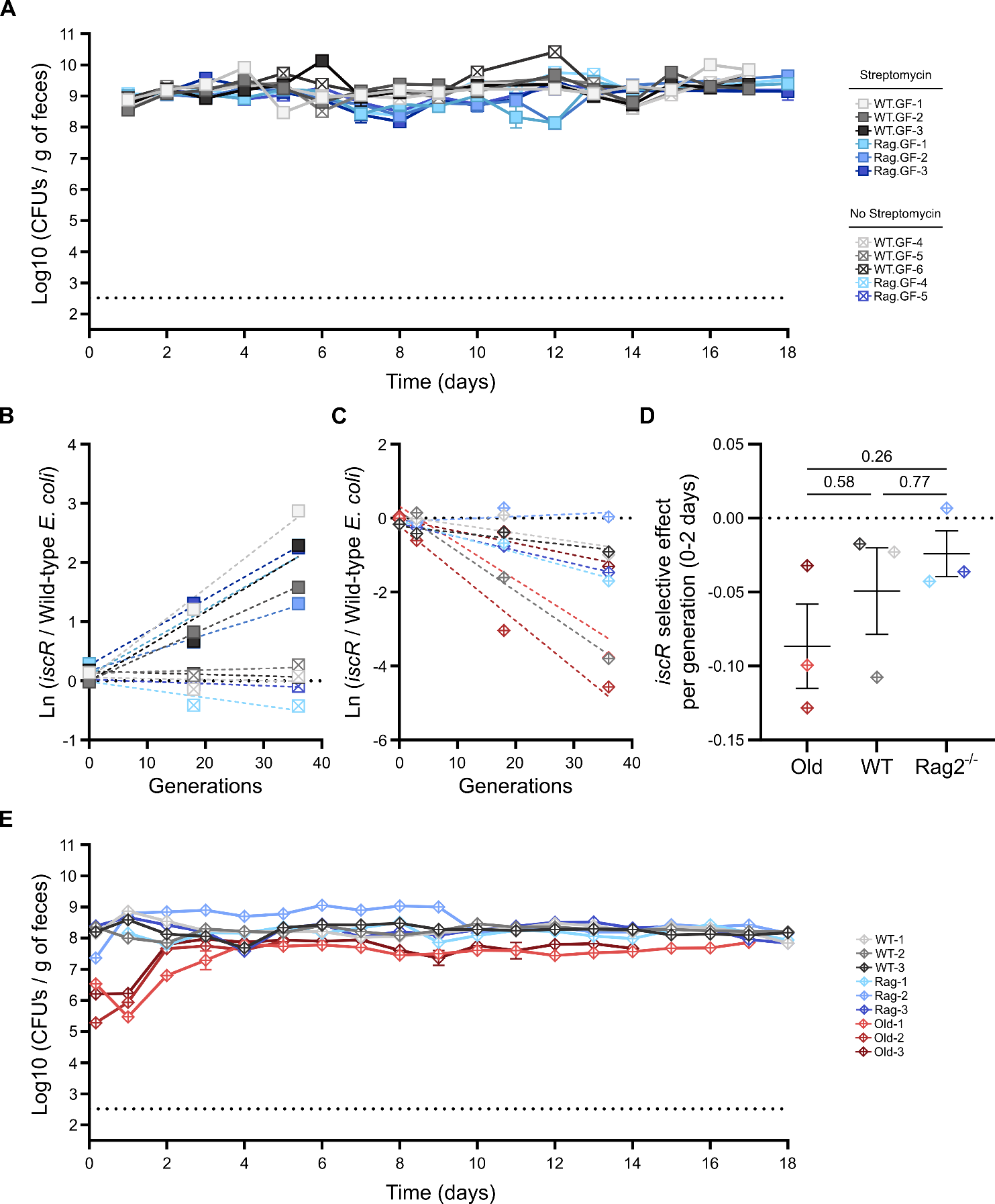

**Figure S2. *E. coli* total loads are independent of the streptomycin treatment and host physiological state*.* Related to Figure 3. (A)** *E. coli* loads per gram of feces throughout the competitions of *iscR* against the wild-type *E. coli* in WT and *Rag2^-/-^* germ-free mice with (WT, n = 3; *Rag2^-/-^*, n = 3) and without (WT, n = 3; *Rag2^-/-^*, n = 2) continuous streptomycin treatment. Error bars represent the ± 2*SEM. The dashed lines indicate the limit of detection; **(B)** The selective effect of *iscR* after two days of competition in WT and *Rag2^-/-^* germ-free mice with (WT, n = 3; *Rag2^-/-^*, n = 3) and without (WT, n = 3; *Rag2^-/-^*, n = 2) continuous streptomycin treatment is inferred from the slope of the linear regression, corresponding to the selection coefficient. The dashed lines represent the best-fit linear regression; **(C)** The selective effect of *iscR* after two days of competition in Old (n = 3), WT (n = 3), and *Rag2^-/-^* (n = 3) mice in the absence of streptomycin treatment is inferred from the slope of the linear regression, corresponding to the selection coefficient. The dashed lines represent the best-fit linear regression; **(D)** Selective effect of *iscR* after two days of competition in Old (-0.09 ± 0.03, n = 3), WT (-0.05 ± 0.03, n = 3) and *Rag2^-/-^* (-0.02 ± 0.02, n = 3) mice in the absence of streptomycin treatment. The middle line indicates the mean, the error bars represent ± SEM. For statistical significance the ANOVA with Tukey’s multiple comparison test was used **(E)** *E. coli* loads per gram of feces throughout the competitions of *iscR* against the wild-type *E. coli* in Old (n = 3), WT (n = 3), and *Rag2^-/-^* (n = 3) mice in the absence of streptomycin treatment. Error bars represent the ± 2*SEM. The dashed lines indicate the limit of detection. See also Table S1.

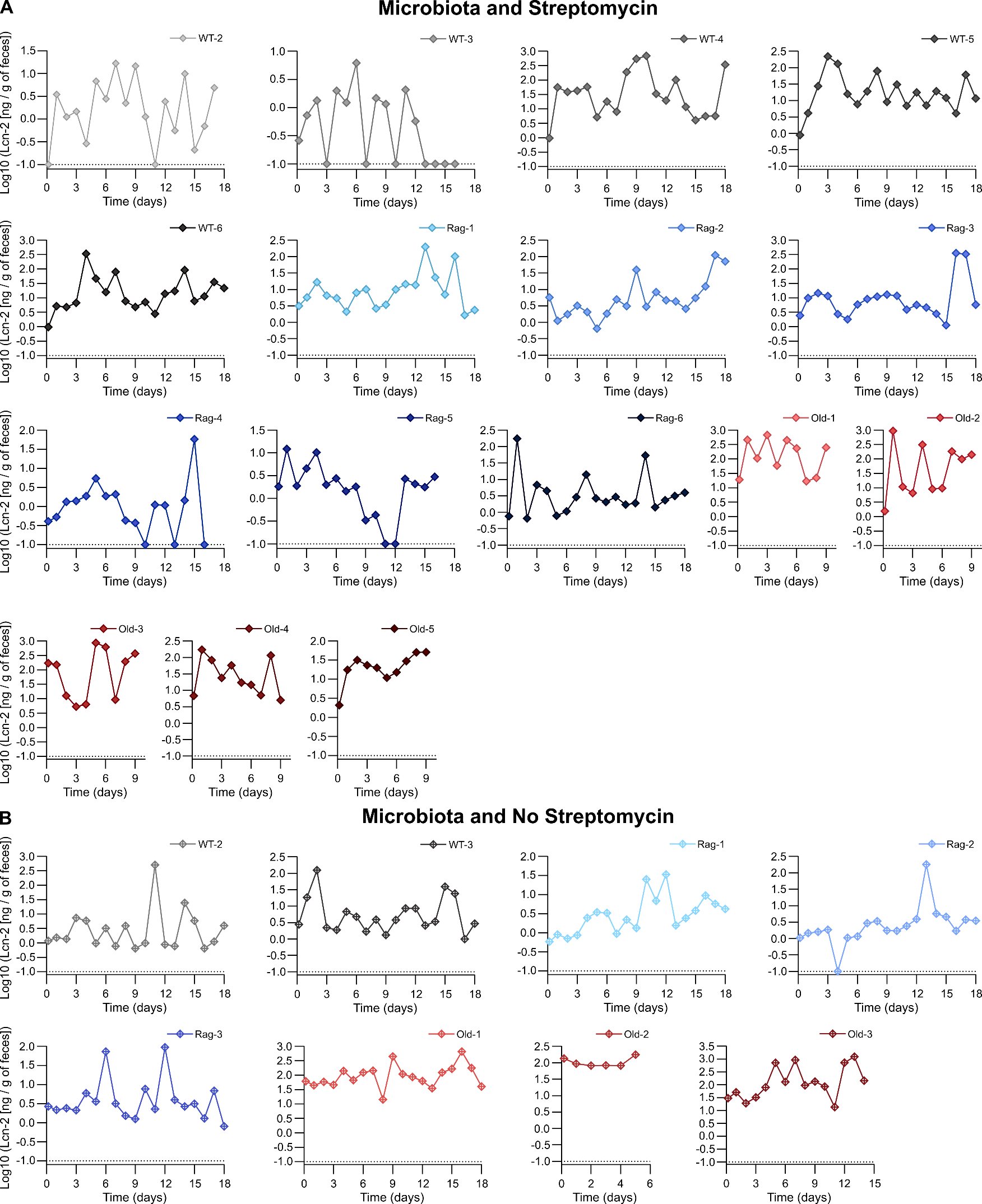

**Figure S3. Variation in fecal Lipocalin-2 in mice with microbiota. Related to Figure 5.** Dynamics of the concentration of fecal Lipocalin-2 during the *in vivo* competitions of *iscR* against the wild-type *E. coli* in mice with microbiota and in the **(A)** presence or **(B)** absence of streptomycin. The dashed lines indicate the limit of detection. See also Table S1.

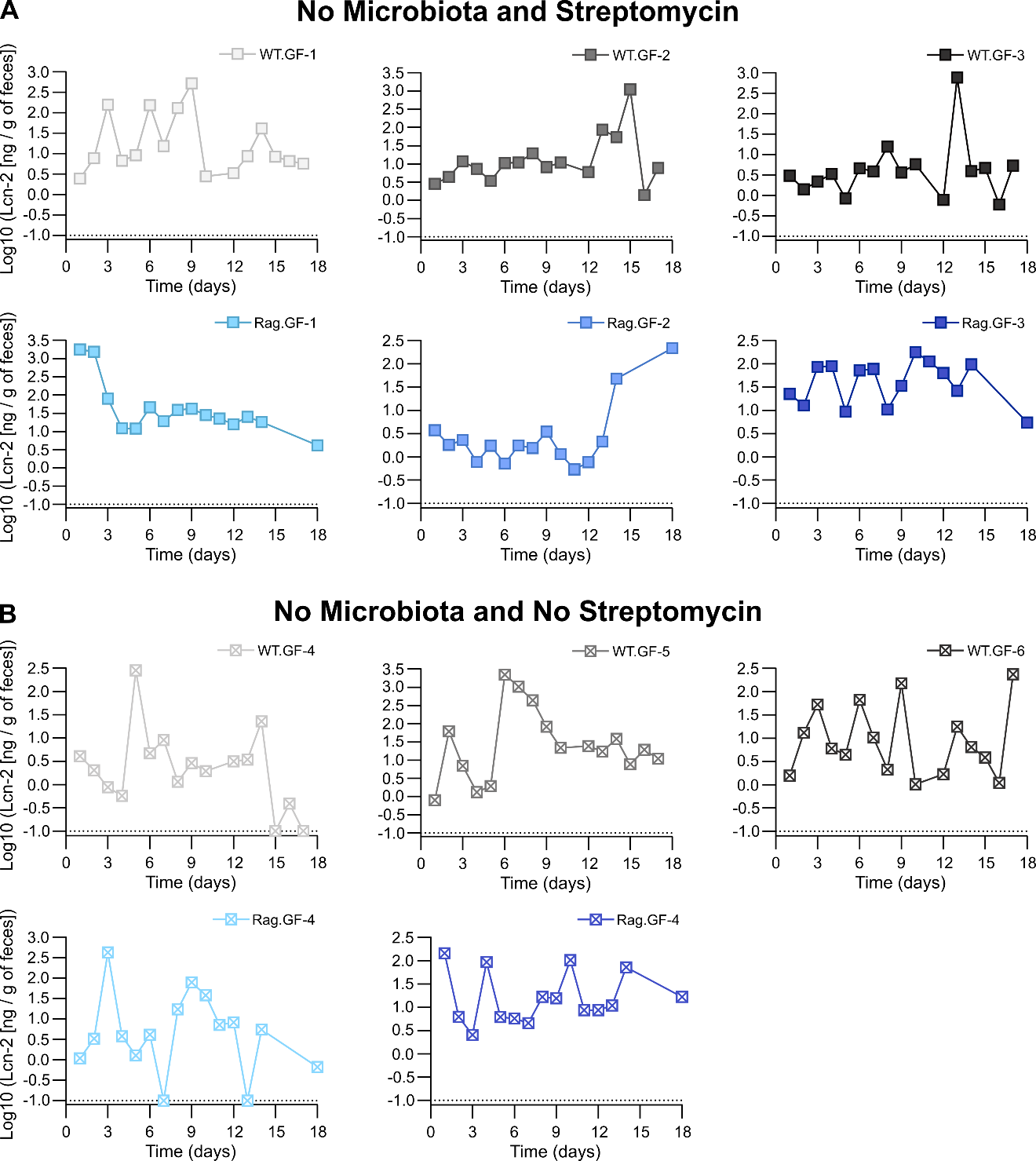

**Figure S4. Variation in fecal Lipocalin-2 in germ-free mice. Related to Figure 5.** Dynamics of the concentration of fecal Lipocalin-2 during the *in vivo* competitions of *iscR* against the wild-type *E. coli* in germ-free mice in the **(A)** presence or **(B)** absence of streptomycin. The dashed lines indicate the limit of detection. See also Table S1.

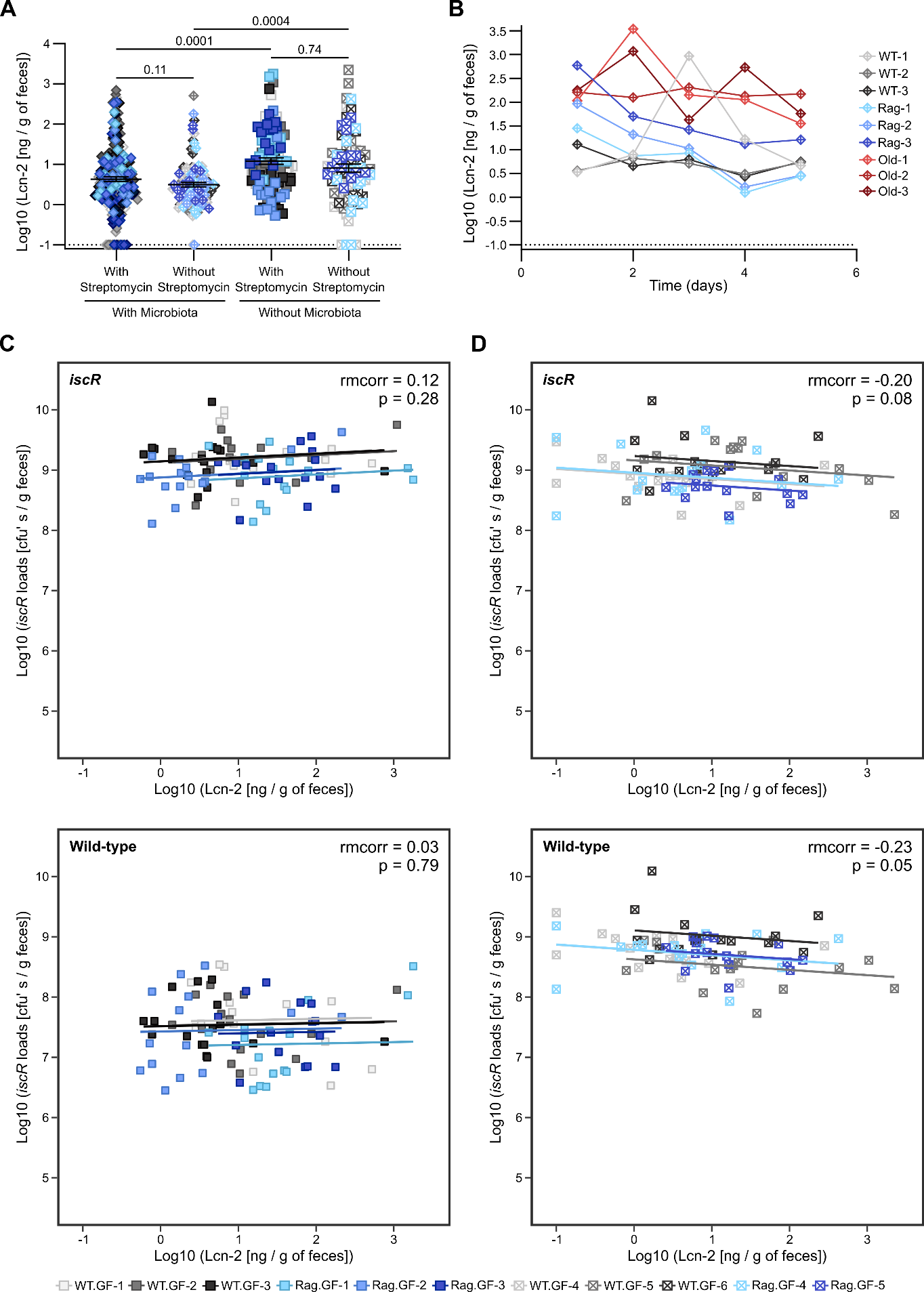

**Figure S5. Variation in fecal Lipocalin-2 levels for manipulated and unmanipulated mice and fecal Lipocalin-2 levels correlation with *E. coli* loads in germ-free mice. Related to Figure 5. (A)** Variation in fecal Lipocalin-2 concentration measured in mice with microbiota in the presence (n = 17) or absence (n = 9) of streptomycin treatment, and germ-free mice in the presence (n = 6) or absence (n = 5) of streptomycin treatment. The dashed lines indicate the limit of detection; **(B)** Dynamics of the concentration of fecal Lipocalin-2 in unmanipulated mice with microbiota (WT, n = 3; *Rag2^-/-^*, n = 3; Old, n = 3). The dashed lines indicate the limit of detection; **(C)** Repeated measures correlation between *iscR* (top panel) or wild-type *E. coli* (bottom panel) abundance and fecal Lipocalin-2 levels in germ-free mice treated with streptomycin (n = 6); **(D)** Repeated measures correlation between *iscR* (top panel) or wild-type *E. coli* (bottom panel) abundance and fecal Lipocalin-2 levels in germ-free mice not treated with streptomycin (n = 5). In panel A the middle line indicates the mean, the error bars represent the ± SEM, and a Kruskal-Wallis test with Dunn’s multiple comparisons test was used. See also Table S1.
